# The environmental stress response controls the biophysical properties of the cytoplasm and is critical for survival in quiescence

**DOI:** 10.1101/2025.09.30.679535

**Authors:** Lorena Kronig, Carmen A. Weber, Pablo Aurelio Gómez-García, Jonas S. Fischer, Christian Doerig, Agnès Michel, Paolo Ronchi, Sarah Khawaja, Paola Picotti, Benoît Kornmann, Karsten Weis

**Affiliations:** Department of Biology, Institute of Biochemistry, ETH Zurich, Zurich, Switzerland; Bringing Materials to Life Initiative, ETH Zurich, Zurich, Switzerland; Department of Biology, Institute of Molecular Systems Biology, ETH Zurich, Zurich, Switzerland; Department of Biochemistry, University of Oxford, Oxford OX1 3QU, UK; EMBL EM Core Facility, European Molecular Biology Laboratory (EMBL), Heidelberg, Germany

**Author notes:** These authors contributed equally (Lorena Kronig, Carmen A. Weber).

## Abstract

All organisms have evolved survival strategies to cope with changes in environmental conditions. Nutrient deprivation, one of the most frequently encountered stresses in nature, causes haploid budding yeast to enter a reversible state of non-proliferation known as quiescence, which entails extensive remodeling of gene expression, metabolism and the cellular biophysical properties. Yeast cells can adapt to and survive long periods of time in glucose starvation-induced quiescence, provided they are able to respire in the early stages of glucose withdrawal. When respiration is blocked during glucose withdrawal, cells prematurely age and exhibit markedly reduced survival and cytoplasmic diffusion. We find here that respiration is required to induce a quiescence-related gene expression program. Induction of this program prior to withdrawing glucose in respiration-inhibited cells bypasses the need for respiration and rescues survival and biophysical properties to levels seen in glucose-starved but respiration-competent cells. This rescue effect relies on proteomic adaptation, which partially occurs through inactivation of Ras/PKA signaling and activation of the environmental stress response via the transcription factors Msn2/4. This signaling cascade triggers the expression of stress response genes and modulates the cytoplasmic diffusion state of cells, ensuring long-term survival in quiescence even in the absence of respiration. Our results highlight the importance of stress adaptation in quiescence and aging, integrating gene expression control and modulation of cytoplasmic properties to maintain cell fitness.

## Introduction

Many cell types can exist for extended periods in a state of non-proliferation known as quiescence (Laporte and Sagot, 2025). Quiescent cells are defined by their ability to re-enter the cell cycle, distinguishing them from senescent and terminally differentiated cells, which have permanently lost the ability to divide (Marescal and Cheeseman, 2020; Opalek et al., 2023). Over time, quiescent cells can gradually lose their ability to re-enter the cell cycle, a process also known as chronological aging (Fabrizio and Longo, 2003). In unicellular eukaryotes, quiescence ensures long-term survival in conditions that do not allow for proliferation, whereas in higher eukaryotes, quiescent cells play diverse and important roles that include tissue regeneration, wound healing and reproduction (Özgüldez and Bulut-Karslioglu, 2024). Entry into and exit from quiescence is carefully regulated, and defects in either can cause or aggravate pathologies such as cancer and auto-immune diseases (Marescal and Cheeseman, 2020). In contrast to the complex signaling governing quiescence in multicellular organisms, unicellular organisms such as budding yeast can transition between proliferation and non-proliferation solely based on nutrient availability, making them useful models for studying this phenomenon (Sun and Gresham, 2021; Laporte and Sagot, 2025). In budding yeast, acute glucose withdrawal or growth into stationary phase causes cells to enter a quiescent state characterized by growth arrest and increased stress resistance (Werner-Washburne et al., 1993; Herman, 2002; Fabrizio and Longo, 2003; Gray et al., 2004; Wood et al., 2020). This transition requires a plethora of cellular adaptations to occur in a coordinated manner.

A striking aspect of this adaptation is a change in the biophysical properties of the cell cytoplasm, which are thought to be actively regulated and relevant for cell survival (Joyner et al., 2016; Munder et al., 2016). The reorganization of the cytoplasm leads to a remarkable increase in both cytoplasmic rigidity and crowding (Joyner et al., 2016; Munder et al., 2016; Heimlicher et al., 2019). These adaptations arise from cell volume reduction and vacuole expansion without a change in total biomass (Joyner et al., 2016), and similar changes have been documented across different organisms and probes (Parry et al., 2014; Joyner et al., 2016; Munder et al., 2016; Heimlicher et al., 2019; Sakai et al., 2024). The regulation of the biophysical properties in quiescent cells is expected to directly affect many cellular processes, including diffusion-limited reactions, intracellular transport, translation, protein folding and aggregation (Mourão et al., 2014; Weiss, 2014; Kuznetsova et al., 2015), potentially impacting signaling and metabolism in a manner critical for transitioning into a non-proliferative, dormant state (Parry et al., 2014; Joyner et al., 2016; Munder et al., 2016; Sakai et al., 2024).

In addition to these biophysical effects, significant metabolic changes occur following quiescence onset. Shortly after glucose deprivation, cells switch from fermentation to respiration, using the fermentation by-product ethanol or internal amino acids and lipid stores as an energy source (Seo et al., 2017; Weber et al., 2020). This metabolic reprogramming allows them to maintain ATP production, albeit at lower levels (Xu and Bretscher, 2014; Joyner et al., 2016; Takaine et al., 2019), and is essential for survival both in acute glucose deprivation (Weber et al., 2020) and in stationary phase (Bonawitz et al., 2007; Aerts et al., 2009; Ocampo et al., 2012). Importantly, respiration-deficient cells display not only decreased survival in glucose starvation but also an exaggerated reduction in cytoplasmic diffusion (Joyner et al., 2016; Munder et al., 2016), suggesting a close link between cytoplasmic diffusion and viability.

Glucose starvation also induces large-scale changes in gene expression, including the activation of the environmental stress response (ESR) and the repression of many growth-promoting ‘house-keeping’ genes (Gasch et al., 2000; Causton et al., 2001). The ESR is induced by the master transcriptional regulators Msn2/4 and coordinates the activation of hundreds of target genes (Gasch et al., 2000; Causton et al., 2001). Msn2/4 activation is regulated via several parallel and interconnected pathways, such as the Ras/cAMP/PKA pathway and the stress-responsive kinases Rim15, Yak1 and Snf1 (Santangelo, 2006; de Virgilio, 2012). In addition to glucose starvation, the ESR is activated upon a variety of other stress triggers, including oxidative stress, osmotic shock, high ethanol, and temperature changes, providing both short-term and long-term stress protection (Gasch et al., 2000; Causton et al., 2001; Berry and Gasch, 2008).

Since the transcription and translation of stress-induced genes require energy, we hypothesized that the respiration-requirement for survival after glucose deprivation reflects at least in part the need to generate sufficient ATP to produce essential quiescence factors. Indeed, we find that inducing a stress response prior to glucose withdrawal bypasses the need for respiration during glucose withdrawal. We show that a preparatory heat shock rescues not only survival but also the biophysical properties of the cytoplasm, restoring the phenotypes of respiration-inhibited, glucose-starved cells to those resembling respiration-competent, glucose-starved cells. The rescue effect requires protein synthesis, is partially dependent on Msn2/4, and can be reproduced by deleting components of the Ras/cAMP/PKA pathway that inhibit Msn2/4. Overall, our results suggest that stress response activation is crucial for proper regulation of the cell’s biophysical properties and for ensuring survival during quiescence.

## Results

### A preparatory heat shock increases survival and rescues cytoplasmic diffusion in respiration-deficient glucose starvation

Respiration-deficient glucose starvation is characterized by a drastic viability defect (Weber et al., 2020) and a severe drop in cytoplasmic diffusion (Joyner et al., 2016; Munder et al., 2016). These concurrent phenotypes prompted us to investigate whether cytoplasmic diffusion is functionally connected to cell survival. First, we confirmed that during glucose starvation, respiration inhibition with Antimycin A (AntA) or by deleting *CBP2* strongly reduced cell survival (**Figures 1A, 1B and S1A**). We hypothesized that the lack of ATP production prevents the induction of protective quiescence genes, underlying the survival defect in respiration-deficient starvation. To test this, we applied a heat shock (HS) and recovery period to glucose-growing cells prior to Antimycin A treatment and glucose withdrawal (**Figure 1A**). HS is known to trigger the rapid induction of both the heat shock response (HSR) and environmental stress response (ESR), resulting in the expression of hundreds of stress-related genes (Gasch et al., 2000; Morano et al., 2012; Verghese et al., 2012) and providing stress tolerance for subsequent stresses (Pereira et al., 2001; Berry and Gasch, 2008). Intriguingly, the viability decrease by respiration inhibition was largely rescued by this preparatory HS (**Figure 1B**), suggesting that the pre-HS indeed prepares the cells for the subsequent respiration-deficient glucose starvation. Importantly, the pre-HS did not rescue the critically low ATP levels in respiration-inhibited glucose starvation (**Figure 1C**).

**Figure 1:**
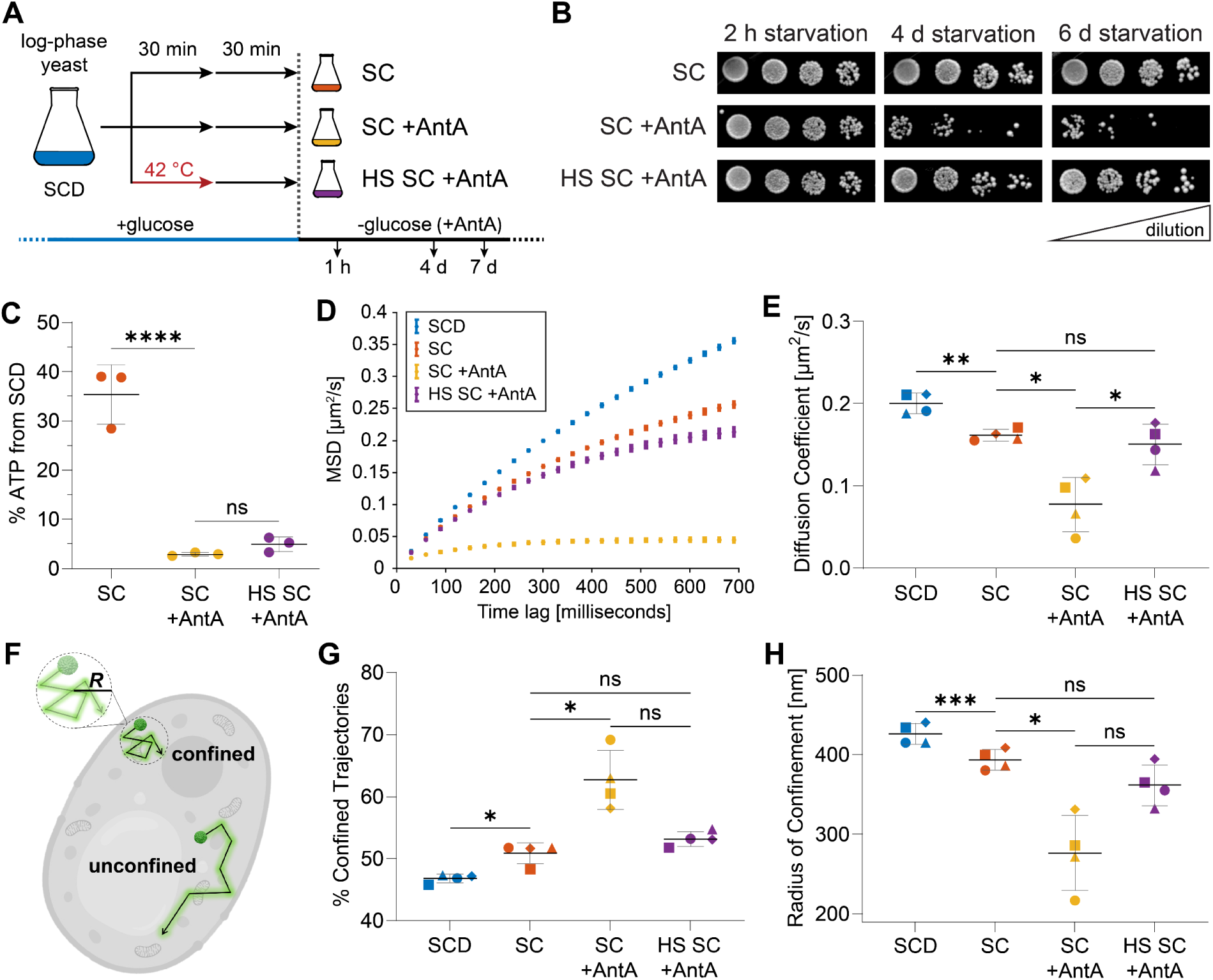
A preparatory heat shock increases survival and rescues cytoplasmic diffusion in respiration-deficient glucose starvation. **A**) Experimental set-up: Yeast cells were grown to logarithmic phase in synthetic complete medium containing glucose (SCD) and washed into glucose-free medium (SC), washed into glucose-free medium with Antimycin A (SC +AntA) or treated with a 42 °C heat shock (HS) and a recovery from the heat shock and then washed into glucose-free medium with Antimycin A (HS SC +AntA). All samples were incubated at 30 °C. **B)** Cells treated as indicated and starved for the indicated time were spotted onto YPD plates. Regrowth ability was used to qualitatively assess survival. **C)** ATP levels were measured after 1 hour treatment in SC, SC +AntA or HS SC +AntA and are shown relative to the ATP level measured in cells growing in SCD (n = 3 independent replicates, mean±SD). One-way ANOVA with Tukey’s multiple comparisons test (****p ≤ 0.0001, ^ns^p > 0.05). **D)** Samples were generated as in A) and starved for 30 minutes. TE-MSD curves show the mean square displacements of all individual trajectories and the S.E.M. The TE-MSD curves shown are from one replicate. **E)** Diffusion coefficients were calculated based on the TE-MSD curves of all trajectories (n = 4 independent replicates, mean±SD). One-way ANOVA with Tukey’s multiple comparisons test (^ns^p > 0.05, *p ≤ 0.05, **p ≤ 0.01). **F)** Illustration of GEMs moving inside the cytoplasm in an unconfined or confined manner. For confined trajectories we determined the radius of confinement (R), estimating the size of the physical space to which the particle is confined. **G)** Percentages of confined trajectories were calculated based on the TE-MSD curves of all trajectories (n = 4 independent replicates, mean±SD). One-way ANOVA with Tukey’s multiple comparisons test (^ns^p > 0.05, *p ≤ 0.05). **H)** Radii of confinement were calculated based on the TE-MSD curves of the confined trajectories (n = 4 independent replicates, mean±SD). One-way ANOVA with Tukey’s multiple comparisons test (^ns^p > 0.05, *p ≤ 0.05, ***p ≤ 0.001).

We next asked if changes in the biophysical properties of the cytoplasm correlate with survival in the aforementioned stress conditions. To probe the dynamics of the cytoplasm, we performed single particle tracking (SPT) using the well-established <u>g</u>enetically <u>e</u>ncoded <u>m</u>ultimeric nanoparticles, or GEMs, which with a size of 40 nm are in the same size range as ribosomes (Delarue et al., 2018). Analyzing the trajectories of GEM particles in hundreds of cells allowed us to obtain the Time-Ensemble Mean Square Displacement (TE-MSD) curve, which in turn was used to calculate the diffusion coefficient (see Methods). The TE-MSD curves (**Figure 1D**) and diffusion coefficients (**Figure 1E**) confirmed the previously reported diffusion reduction in glucose-starved cells (−19%, *SC*) compared to logarithmically growing cells (*SCD*), as well as the more pronounced diffusion drop upon additional respiration inhibition (−61%, *SC +AntA*). Remarkably, the pre-HS mitigated the drastic diffusion reduction in respiration-inhibited glucose starvation (−25%, *HS SC +AntA*) even though the ATP levels do not recover.

To further characterize the movement of GEMs, we classified the trajectories into two motion types, confined or unconfined, based on their anomalous exponents (**Figure 1F**, see Methods) and determined the proportion of these two groups in the different conditions. In *SCD*, 47% of trajectories were confined, in *SC* this proportion was increased to 51%, in *SC +AntA* to 63% and in *HS SC +AntA* to 53% (**Figure 1G**). The radius of confinement, which measures the physical space to which particles are confined (*R*, see Methods), exhibits the same trend as the percentage of confined trajectories; it was reduced by 8% in *SC*, by 35% in *SC +AntA* and by 15% in *HS SC +AntA* (**Figure 1H**). Thus, lower diffusion is accompanied by a higher proportion of confined trajectories and a decrease in the size of the confined spaces in which the particles diffuse. Together, our results show that the HS-mediated improvement of cell survival in respiration-inhibited glucose starvation is accompanied by a rescue in cytoplasmic diffusion, indicating that HS-triggered cytoplasmic adaptation enhances tolerance for subsequent nutrient stress and entry into quiescence.

### Macromolecular crowding-induced confinement causes cytoplasmic reorganization in quiescence

To understand the biophysical changes underlying the reduced diffusion in quiescent cells, we examined how cytoplasmic organization and crowding are altered in our quiescence conditions. We used FIB-SEM 3D imaging on high-pressure-frozen non-starved (*SCD*) and 30-minute glucose-starved (*SC*) cells to examine the macromolecular density in the cytoplasm. Determining the concentration of ribosomes, which are readily detectable as electron-dense particles, showed a 28% density increase in glucose-starved cells compared to non-starved cells (**Figures 2A and 2B**). The FIB-SEM images also revealed an increase in vacuolar size (**Figure 2A**), which has been reported previously (Joyner et al., 2016). These results are in line with our previous finding that glucose starvation reduces the cytoplasmic space that is available for diffusing molecules by ∼30%, and the hypothesis that cytoplasmic volume reduction leads to increased crowding, reducing particle mobility and diffusion (Joyner et al., 2016). To validate that crowding can indeed modulate GEM diffusion in yeast cells, we induced higher crowding with hyperosmotic shocks using NaCl. Both cell volume and GEM diffusion declined progressively with increasing NaCl concentration (**Figures 2C, 2D and 2E**). Hyperosmotic shock also led to an increase in the percentage of confined trajectories and to a smaller radius of confinement (**Figures 2F and 2G**).

**Figure 2:**
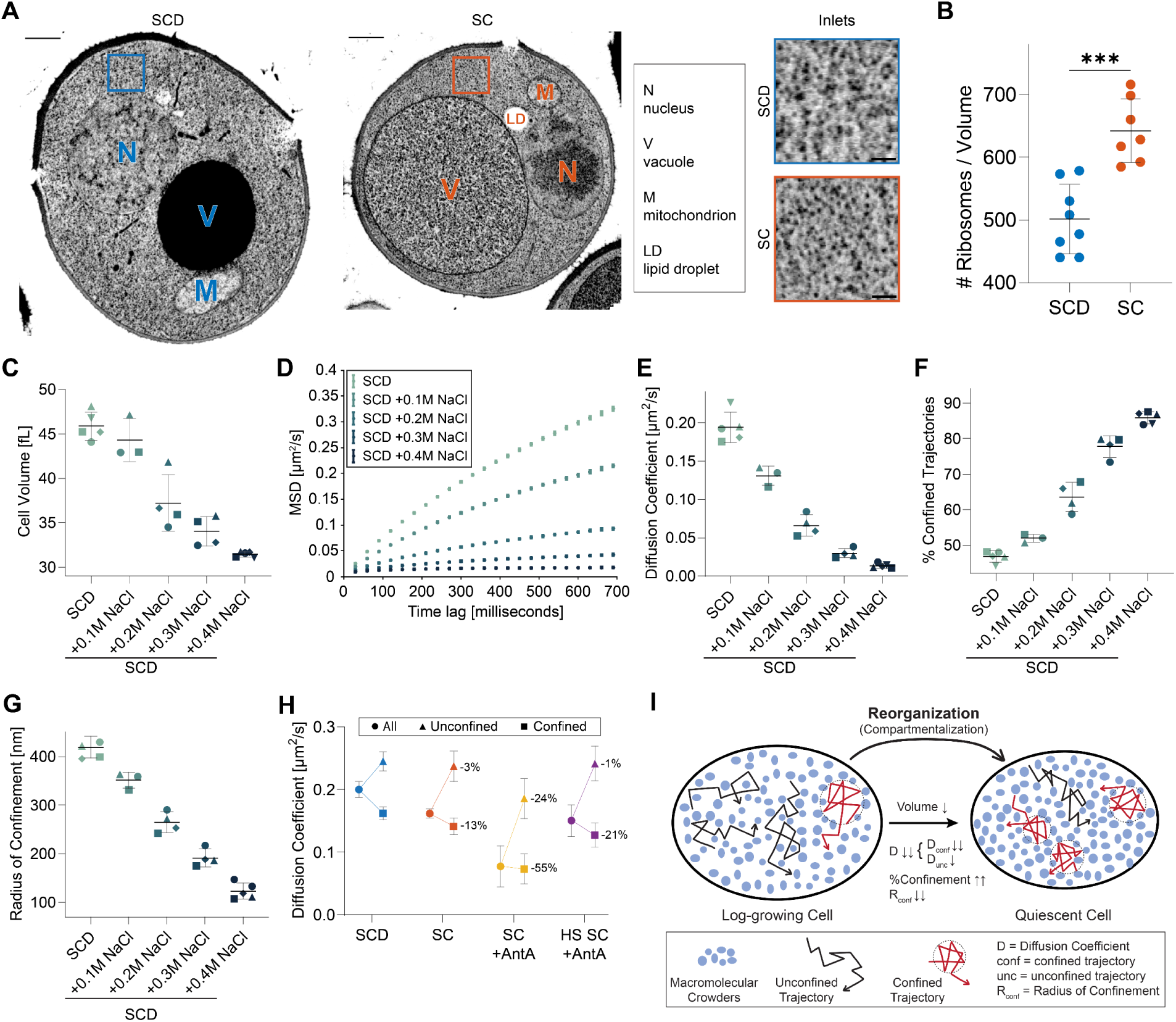
Macromolecular crowding-induced confinement causes cytoplasmic reorganization in quiescence. **A)** Single slices extracted from the 3D volumes imaged by FIB-SEM of non-starved (SCD) and 30-minute glucose-starved (SC) cells. Full image scale bar = 500 nm, inlet scale bar = 100 nm. **B)** Quantification of macromolecules within equal volumes of the bulk cytoplasm in non-starved and starved cells imaged with FIB-SEM. 3-4 volumes in 2 different cells per condition were analyzed blindly (mean±SD). Unpaired two-tailed t-test (***p-value = 0.0002). **C)** Cell volume measurements of cells treated with increasing NaCl concentrations for 5 minutes (n ≥ 4 independent replicates, mean±SD). **D)** Time-Ensemble Mean Square Displacement (TE-MSD) curves of all trajectories observed in cells grown in glucose medium and treated with increasing amounts of NaCl for 5 minutes. The TE-MSD curves show the mean square displacements of all individual trajectories and the S.E.M. The TE-MSD curves shown are from one replicate. **E)** Diffusion coefficients were calculated based on the TE-MSD curves of all trajectories (n ≥ 4 independent replicates). **F)** Percentages of confined trajectories were calculated based on the TE-MSD curves of all trajectories (n ≥ 4 independent replicates, mean±SD). **G)** Radii of confinement were calculated based on the TE-MSD curves of the confined trajectories (n ≥ 4 independent replicates, mean±SD). **H)** Diffusion coefficients of the distinguishable populations of trajectories. The diffusion coefficients were either calculated from all trajectories (circles), from the unconfined trajectories (triangles) or from the confined trajectories (squares). The percentage changes compare the confined or unconfined trajectories’ diffusion coefficients in each starvation condition with the SCD condition. **I)** Illustration showing the reorganization of the cytoplasm as cells are starved of glucose and enter quiescence.

We next asked if the changes in diffusion of the 40 nm GEM particle are due to uniformly altered crowding or arise from localized confinement due to compartmentalization. To this end, we analyzed the diffusion coefficients of all trajectories, as well as those of confined and unconfined trajectories separately. In all starvation conditions, the diffusion coefficient of the confined trajectories exhibited a more pronounced decrease than the unconfined ones, when compared to non-starved cells (**Figure 2H**). This, together with the increased percentage of confined trajectories (**Figure 1G**), is the main cause of the overall diffusion decrease in quiescence. Therefore, lower diffusion in quiescence is not caused by a uniform increase in crowding or viscosity, but rather by a general reorganization of the cytoplasm, featuring more, smaller and potentially denser compartments in which the GEMs diffuse more slowly (**Figure 2I**). Additional evidence for this comes from the observation that imaging GEMs at a lower frame rate (i.e., longer exposure time) showed a uniformly diffuse signal in *SCD,* whereas distinct foci appeared under starvation conditions (**Figures S2A and S2B**). This might suggest that GEMs aggregate in starvation. However, analyzing GEM particle intensities in fixed cells imaged at faster frame rate (used for SPT) showed uniform GEM brightness distributions in all conditions (**Figure S2C**). Rather than aggregates, the bright foci thus represent immobile, single GEM particles that are confined and appear brighter during long exposure times due to their inability to move. Indeed, the percentage of cells with GEM foci in these conditions followed the same trends as the amount of confinement (**Figure S2B**). This analysis underscores the need for sensitive and rapid imaging techniques to probe cytoplasmic diffusion.

Taken together, these findings show that quiescence as well as hyperosmotic shock lead to a crowding-induced restructuring of the cytoplasm, likely altering diffusion-driven processes and affecting cell function and fitness. The critically low diffusion and high confinement observed in the *SC +AntA* condition is rescued by the pre-HS, presumably allowing cells to adapt to and survive in *SC +AntA*.

### Heat shock-induced proteomic adaptation primes cells for respiration-deficient glucose starvation

To determine if the HS rescue effect on cytoplasmic diffusion and cell survival is mediated by changes in gene expression through stress response activation, we tested whether translation is required during the pre-HS. Translation inhibition using cycloheximide (*CHX*) during the HS and recovery period indeed abrogated the benefit on survival and cytoplasmic diffusion (**Figures 3A and 3B**). Thus, the HS-induced benefit is lost when cells are unable to produce new proteins.

**Figure 3:**
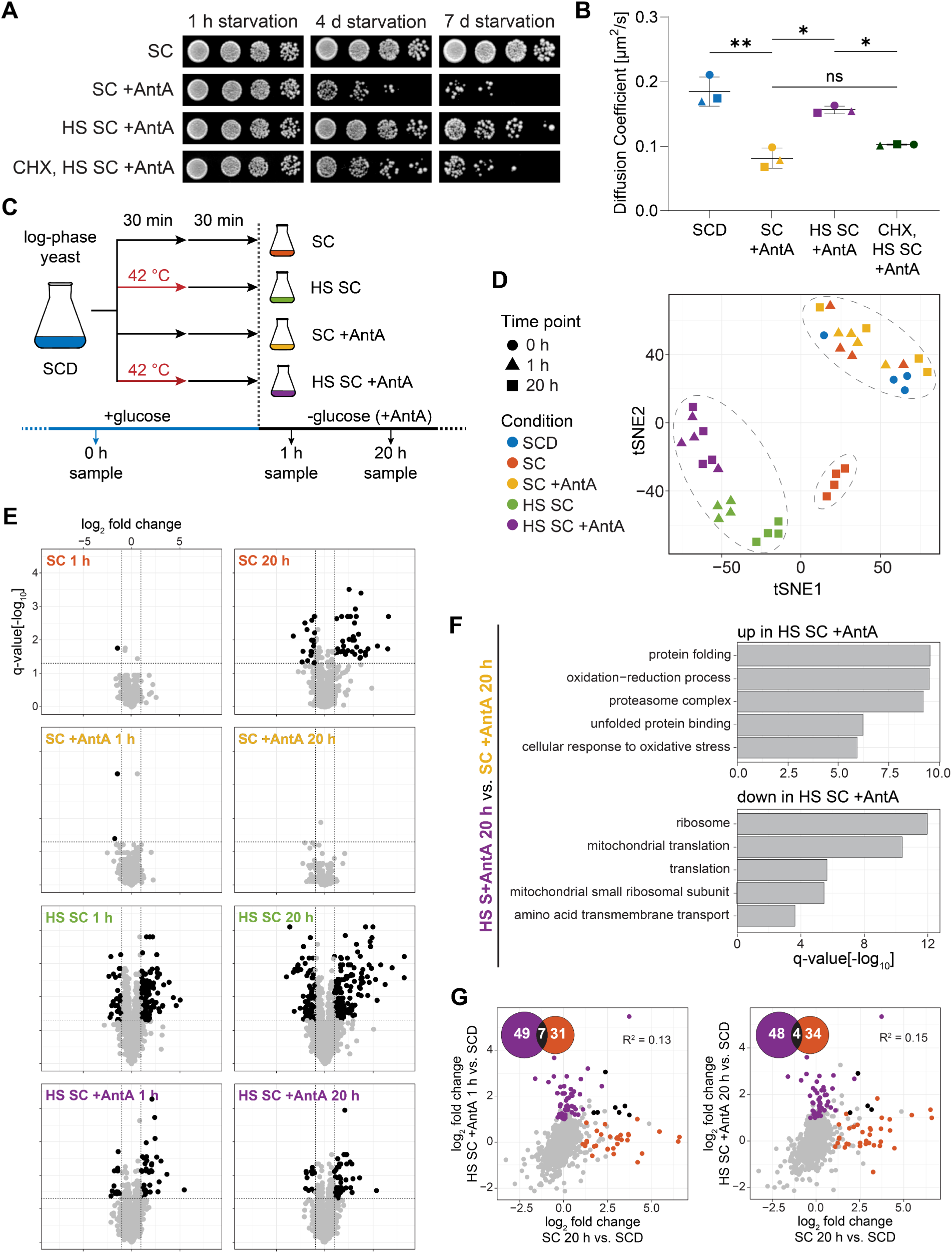
Heat shock-induced proteomic adaptation primes cells for respiration-deficient glucose starvation. **A)** Logarithmically growing yeast cells were treated as described in Figure 1A. One additional sample was treated with cycloheximide (CHX) 5 min prior to the HS, during the HS and recovery and then washed into glucose-free medium with Antimycin A without CHX (CHX, HS SC +AntA). All samples were incubated at 30 °C and cells were spotted onto YPD plates at the indicated time points. Regrowth ability after 2 days was used to qualitatively assess survival. **B)** Samples were generated as in A) and starved for 30 minutes. Diffusion coefficients were calculated based on the TE-MSD curves of all trajectories (n = 3 independent replicates, mean±SD). One-way ANOVA with Tukey’s multiple comparisons test (^ns^p > 0.05, *p ≤ 0.05, **p ≤ 0.01). **C)** Scheme of sample generation: Yeast cells were grown to logarithmic phase in synthetic complete medium containing glucose (SCD) and washed into glucose-free medium (SC), washed into glucose-free medium with Antimycin A (SC +AntA) or treated with a 42 °C heat shock (HS) and a recovery from the heat shock and then washed into glucose-free medium without (HS SC) or with Antimycin A (HS SC +AntA). For the SCD condition, cells were collected in logarithmic phase (0 h). For the other conditions, one sample was collected after 1 hour and one after 20 hours of starvation. The proteomics experiment was done in 4 biological replicates. **D)** t-distributed stochastic neighbor embedding (tSNE) clustering of proteomic samples. Each of the four replicates is shown. **E)** Protein abundance differences between logarithmically growing SCD cells and cells treated with the indicated stress condition. Data points correspond to the log2 fold change of the mean of four replicates, p-values were calculated using a two-tailed Student’s t-test and adjusted for multiple testing using the Benjamini-Hochberg correction. **F)** Gene ontology (GO) analysis comparing 20 h HS SC+AntA and 20 h SC +AntA conditions. GO terms of proteins in the top and bottom 10% of fold changes were tested for significance using a hyper-geometric test and resulting p-values were adjusted for multiple comparisons using the Benjamini-Hochberg correction. **G)** Correlation of protein abundance changes from E), comparing 1 h HS SC +AntA with SCD and 20 h SC with SCD or 20 h HS SC +AntA with SCD and 20 h SC with SCD. The number of significantly upregulated proteins in each comparison are shown in the Venn diagram and colored accordingly in the correlation plot. Coefficients of determination R^2^ = 0.13 and R^2^ = 0.15.

To uncover the protective adaptation behind the HS benefit, we analyzed the proteome of logarithmically growing cells (*SCD*, single time point) and of four quiescence: glucose starvation (*SC*), respiration-deficient glucose starvation (*SC +AntA*), pre-heat shocked glucose starvation (*HS SC*) and pre-heat shocked, respiration-deficient glucose starvation (*HS SC +AntA*) after 1 hour and 20 hours of starvation (**Figure 3C**). Clustering analysis revealed that the replicates are generally clustered, and the conditions broadly form three different groups (**Figure 3D**). Short-term glucose starvation (*SC 1 h*) and respiration-deficient glucose starvation (*SC +AntA 1 h* and *SC +AntA 20 h*) cluster together with the logarithmic growth condition (*SCD*). This indicates that short-term glucose starvation does not trigger a global reshaping of the protein landscape. Furthermore, cells in respiration-deficient glucose starvation are not able to induce any significant proteomic changes, likely due to the absence of energy. Surprisingly, long-term glucose-starvation (*SC 20 h*) clusters separately from the pre-HS conditions (*HS SC* and *HS SC +AntA*), suggesting that the heat treatment leads to a distinct proteome adaptation.

Comparing logarithmic growth (*SCD*) to each quiescence condition, the individual protein changes confirmed that short-term glucose-starved (*SC 1 h*) and respiration-deficient glucose-starved (*SC +AntA 1 h* and *SC +AntA 20 h*) cells exhibit no or only minimal changes in their proteome, while both pre-HS and long-term glucose starvation (*SC 20 h*) induced broad changes (**Figure 3E**). This suggests that the viability defect of *SC +AntA* cells is not due to specific detrimental changes but rather their inability to adapt effectively to the change in growth conditions.

To better characterize which genes are differentially regulated by the pre-HS, we performed a Gene Ontology (GO) analysis comparing the 20-hour time point of *HS SC +AntA* to the 20-hour time point of *SC +AntA*. Enriched in *HS SC +AntA* were GO terms of the classical ESR including protein folding, proteasomal function, proteostasis and the oxidative stress response (Gasch and Werner-Washburne, 2002) (**Figure 3F**). On the other hand, proteins associated with ribosomes, translation and mitochondrial translation are downregulated in *HS SC +AntA* compared to *SC +AntA*.

Furthermore, the proteins that are enriched in long-term starvation (*SC 20 h vs. SCD)* and those enriched in pre-HS respiration-deficient starvation (*HS SC +AntA 1 h* vs. *SCD; HS SC +AntA 20 h* vs. *SCD*) are poorly correlated and there is only a small overlap of factors that are significantly upregulated in both conditions (**Figure 3G**). Looking at the Gene Ontology (GO) terms of the proteomic changes across all conditions, we observed that long-term starvation (*SC 20*) leads to the upregulation of a few groups of proteins, mostly involved in respiratory activity and rewiring of the metabolism (**Figure S3A**). On the other hand, HS induces proteome changes in several processes, including proteasomal activity, protein folding, osmotic and oxidative stress response, as well as ribosome and translation regulation. To test whether proteins that are induced in the *SC 20 h* sample are potentially also upregulated in the *HS SC +AntA* samples albeit to a lesser extent and below the significance threshold, we highlighted those proteins in the short- and long-term *HS SC +AntA* vs. *SCD* plots **(Figure S3B)**. This shows that most proteins that are induced in the *SC 20 h* sample are not enriched in either of the *HS SC +AntA* samples irrespective of the significance cut-off (**Figure S3B**). Similarly, most of the downregulated proteins in the *SC 20 h* sample are not downregulated in 1 hour of *HS SC +AntA*, however, many do become downregulated after 20 hours of *HS SC +AntA*.

These results suggest that the HS triggers a proteomic adaptation which cannot be induced in respiration-deficient glucose starvation. The HS-induced reprogramming is distinct from the response to glucose starvation, but is nonetheless able to compensate for it, allowing cells to persist in quiescence. The most striking group of HS-induced proteins are involved in proteostasis, protein folding and proteasomal processes. Interestingly, the *HS SC +AntA* condition also exhibits reduced ribosomal and translation-associated proteins, suggesting that proteostasis regulation might be a crucial aspect of the HS-mediated benefit.

### The environmental stress response is critical for survival in quiescence

To investigate the underlying molecular mechanisms of the HS-induced survival benefit, we conducted a <u>SA</u>turated <u>T</u>ransposon <u>A</u>nalysis in <u>Y</u>east (SATAY) screen (Michel et al., 2017). Here, a very large library of yeast clones is created through random transposon insertion leading to gene disruptions and allowing for a selection of detrimental or beneficial insertion events. The transposon library was subjected to a primary experimental condition and two control conditions for comparison. The main condition involved a pre-HS and recovery period followed by respiration-inhibited glucose starvation for 4 days (*4 d HS SC +AntA*). The first control was respiration-inhibited glucose starvation for 1 hour (*1 h SC +AntA*) to control for short-term effects of respiration-inhibited glucose starvation. The second control was a pre-HS and recovery with subsequent respiration-inhibited glucose starvation for one hour (*1 h HS SC +AntA*) to control for effects of the heat shock prior to starvation. Clones that are able to grow are collected and sequenced *en masse* to determine which gene loci can be disrupted without impacting survival. Gene loci which are critical for survival will be underrepresented in the transposon insertion library. We then screened for gene disruptions that are specifically enriched or absent in the 4-day sample (*4 d HS SC +AntA*).

The screen revealed several highly significant gene disruptions that promote or impair the HS-mediated rescue of respiration-deficient, glucose-starved cells (**Figure 4A**). The most significant gene hits that impair survival in *4 d HS SC +AntA* when disrupted are involved in a variety of cellular processes, including ribosome biogenesis (e.g., *ARX1*, *BUD22*, *RRP6*), mitochondrial respiration (e.g., *CBR1*, *COX23*, *ETR1*), and ESR (e.g., *MSN2*, *PSR2*, *YAP1*). Notable among them is *MSN2* and, less significantly scored, *MSN4*, the master regulators of the ESR. The most significant gene hits with enriched disruptions, promoting survival in *4 d HS SC +AntA,* are involved in glucose sensing and Ras/PKA signaling (*GPR1, GPA2, RAS1, RAS2, SHR5, CYR1, CDC25, TPK1, TPK3*). *GPA2*, *RAS2* and *GPR1* all respond to the presence of glucose and trigger cAMP production, causing protein kinase A (PKA) activation, which is a negative regulator of the ESR (**Figure 4B**). The catalytic subunit isoforms of PKA, *TPK1* and *TPK3*, are also significantly promoting survival when disrupted. These results suggest that activation of the ESR through the pre-HS confers a survival benefit on respiration-deficient cells during starvation. This pathway is likely already active prior to starvation through disruption of *RAS2, GPR1* or *GPA2*, as well as upon partial loss of PKA function.

**Figure 4:**
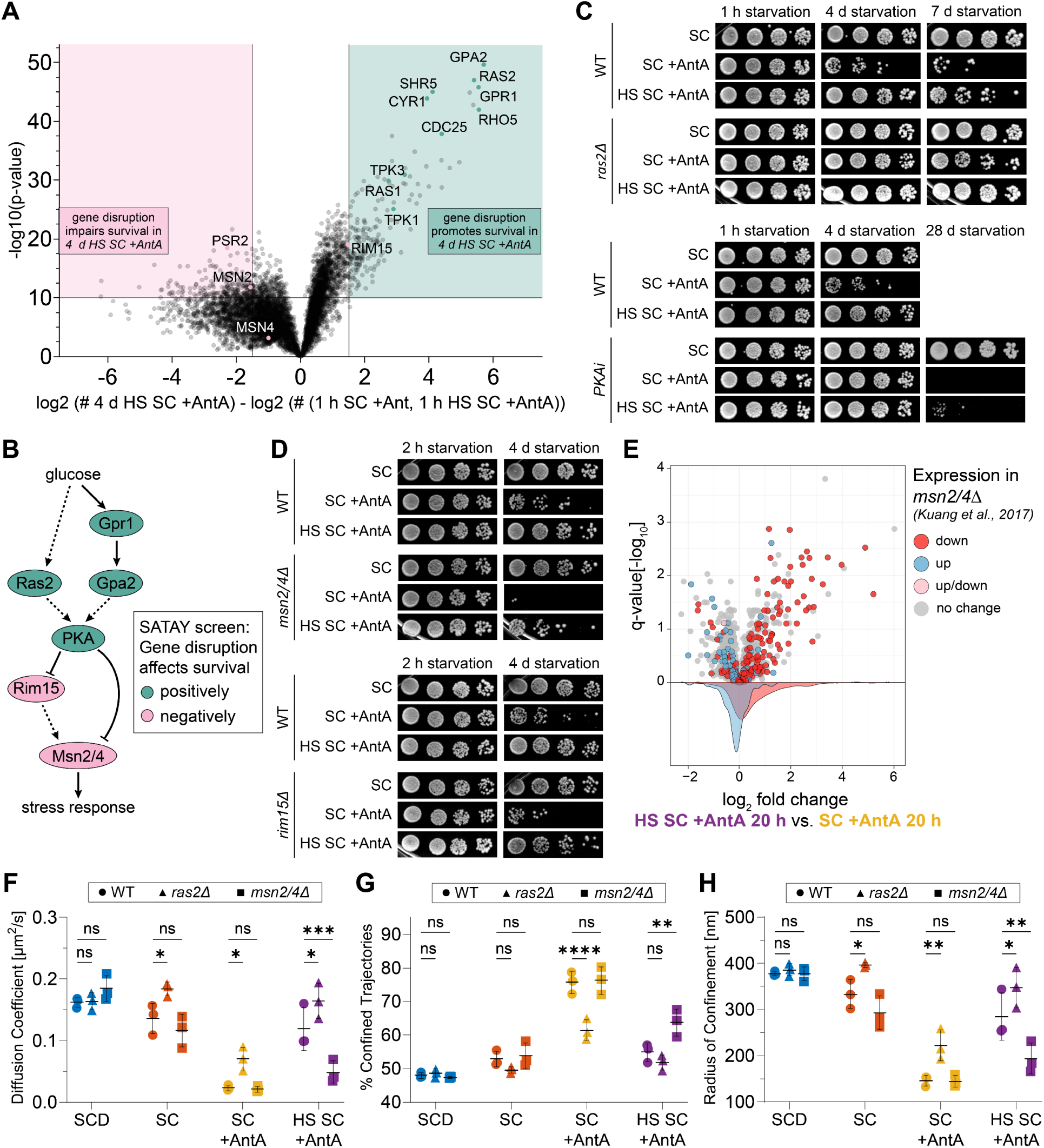
The environmental stress response regulates survival and diffusion in quiescence. **A)** Volcano plot of the SATAY screen results. Positive fold changes (right side of the plot) encompass gene disruptions that led to increased survival in 4 d HS SC +AntA, resulting in the gene sequences being relatively more abundant in the population. Negative fold changes (left side of the plot) encompass gene disruptions that led to decreased survival in 4 d HS SC +AntA, resulting in the gene sequences being relatively less abundant in the population. **B)** Schematic representation of the glucose signaling pathway highlighting the prominent screen hits. Arrows indicate activation (➔) or repression (┫), solid lines indicate direct regulation, dashed lines indicate indirect regulation. **C+D)** Survival assays of wild-type (WT) and mutant cells treated with the indicated conditions and starved for the indicated number of hours or days prior to spotting and re-growth on YPD plates. PKA inhibition (PKAi) was achieved by mutating *TPK1*, *TPK2* and *TPK3* to be sensitive to the ATP-analogue 1NM-PP1, which was added to the logarithmically growing cells for 4 hours prior to washing them into SC or SC +AntA. 1NM-PP1 was also present during the heat shock and recovery, but absent in all conditions after washing. **E)** Protein abundance differences comparing respiration-deficient 20 h glucose starved cells with (HS SC +AntA) or without (SC +AntA) a preparatory heat shock. Data points correspond to the log2 fold change of the mean of four replicates. Unpaired two-tailed t-test with Benjamini-Hochberg correction for multiple testing. The proteins in the volcano plot and the corresponding distributions are colored based on their expression change in the Msn2/4 deletion mutant (Kuang et al., 2017). **F)** Samples were generated as in Figure 1A, using wild-type (WT), ras2Δ, or msn2/4Δ cells. The cells were starved for 1 hour. Diffusion coefficients were calculated based on the TE-MSD curves of all trajectories (n = 3 independent replicates, mean±SD). Two-way ANOVA with Dunnett’s multiple comparisons test (^ns^p > 0.05, **p ≤ 0.01, ****p ≤ 0.0001). **G)** Percentages of confined trajectories were calculated based on the TE-MSD curves of all trajectories (n = 3 independent replicates, mean±SD). Two-way ANOVA with Dunnett’s multiple comparisons test (^ns^p > 0.05, *p ≤ 0.05, **p ≤ 0.01, ****p ≤ 0.0001). **H)** Radii of confinement were calculated based on the TE-MSD curves of the confined trajectories (n = 3 independent replicates, mean±SD). Two-way ANOVA with Dunnett’s multiple comparisons test (^ns^p > 0.05, *p ≤ 0.05, **p ≤ 0.01).

To validate the results of the SATAY screen and confirm our hypothesis, we performed survival assays with a selection of mutants of interest. We observed that deletion of *RAS2, GPR1* or *GPA2* indeed provided a survival benefit in respiration-inhibited glucose starvation, and an even stronger benefit was observed upon inhibition of PKA (**Figures 4C and S4A**). Since deletion of the PKA catalytic subunit isoforms *TPK1*, *TPK2* and *TPK3* causes cell growth arrest, we used a strain with all *TPK* isoforms mutated to be ATP-analogue sensitive to inhibit PKA (*PKAi*) in a controlled manner (**Figure 4C**). Notably, a pre-HS in each of these mutants tested still improved survival compared to *SC +AntA*, suggesting that part of the beneficial, HS-induced response is independent of the Ras/PKA signaling pathway.

Among the detrimental gene disruptions, *MSN2/4* were of particular interest, as they are negatively regulated by PKA and the primary transcriptional ESR activators. In line with the model that ESR activation is critical for the HS benefit, the double deletion of *MSN2* and *MSN4* greatly impaired the survival of respiration-deficient cells in glucose starvation (**Figure 4D**). Interestingly, *RIM15*, one of the upstream activators of Msn2/4, is enriched in transposons, suggesting that its disruption promotes survival in *4 d HS SC +AntA*. However, most of the transposon insertions in *RIM15* were targeting its N-terminal PAS domain, which is thought to act as a regulator of the protein, potentially leading to its constitutive activation and thereby a positive effect on survival (Swinnen et al., 2006). Consistent with this, overexpressing a constitutively active allele of *RIM15,* named *RIM15-S5A* (Reinders et al., 1998), provided a survival benefit (**Figure S4B**) whereas deletion of *RIM15* indeed impaired survival (**Figure 4D**), although to a lesser extent than double-deletion of *MSN2/4*. In both *msn2/4Δ* and *rim15Δ* deletion mutants, the pre-HS treatment improved survival in *SC +AntA,* again suggesting that the Msn2/4-regulated stress response is not the only HS-induced pathway providing a survival benefit. Consistent with this, the strong long-term benefit of PKA inhibition was reduced with additional *MSN2/4* deletion, but these cells nonetheless exhibited improved survival compared to WT or *msn2/4Δ* cells (**Figure S4C**), further hinting at the presence of an Msn2/4-independent benefit of PKA inhibition.

In summary, the SATAY screen shows that inhibiting the glucose sensing pathway is beneficial for survival in respiration-inhibited glucose starvation. We observe a clear survival benefit by disrupting Ras/PKA signaling, and a defect by deleting the ESR activators Msn2/4. However, neither of these disruptions fully abrogate the HS-mediated improvement, suggesting that there are additional, parallel pathways that can confer starvation resistance.

Somewhat surprisingly, the crucial survival genes identified in the SATAY screen are not significantly up- or downregulated in the proteome. More generally, there is a poor correlation of proteins that are abundant in the *HS SC +AntA 20 h* condition and the genes identified as crucial for survival in *HS SC +AntA* in the SATAY screen (**Figure S5A**). The lack of overlapping genes and proteins could be due to functionally redundant Msn2/4 ESR targets that are not detectable in the SATAY screen but are differentially expressed and detected in the proteomics data set.

We therefore also investigated positively regulated Msn2/4 targets, based on published RNA-Seq data (Kuang et al., 2017) (expression down in *msn2/4Δ*). We found that the majority of the genes that are induced by Msn2/4 have upregulated protein levels in the HS conditions (**Figure S3B**), with stronger expression upregulation in *HS SC* than in *HS SC +AntA*. On the other hand, after 20 hours of glucose starvation, only a small subset of Msn2/4-activated targets are translationally upregulated, and they largely do not overlap with *HS SC +AntA 20 h* (**Figure S3B**). Rather, the 20 hours of glucose starvation seems to induce a specific set of Msn2/4 target proteins, which is distinct from the heat-induced response.

Focusing on the proteomes of the 20-hour *SC +AntA* and *HS SC +AntA* conditions, **Figure 4E** highlights genes that are differentially expressed (up, down, no change) in unstressed *msn2/4Δ* cells compared to the wild type (WT). Among the proteins that are enriched in *HS SC +AntA*, we find many targets that are induced by Msn2/4, including genes involved in protein folding and degradation and oxidative stress. On the other hand, many negatively-regulated Msn2/4 targets are less abundant in the *HS SC +AntA* condition (**Figure 4E**). This indicates that HS-triggered Msn2/4 activation leads to the long-lasting upregulation of downstream targets, allowing cells to survive in respiration-deficient glucose starvation. Our survival experiments confirm this, while also revealing beneficial effects of a pre-HS or PKA inhibition that are Msn2/4-independent.

### The environmental stress response regulates diffusion in quiescence

Since the *ras2Δ* and *msn2/4Δ* mutants displayed pronounced positive and negative survival phenotypes, respectively, we also investigated these mutants for changes in cytoplasmic diffusion. In glucose-rich medium (*SCD*), the diffusion behavior (diffusion coefficient, percentage of confined trajectories, radius of confinement) of the mutants is comparable to the wild type (**Figure 4F–H**). The *ras2Δ* mutant exhibited a higher diffusion coefficient and reduced confinement in all starvation conditions, suggesting that Ras/PKA signaling disruption provides an adaptive benefit in the cytoplasmic properties, which correlates with the survival benefit. Notably, we still observed a HS-mediated increase in diffusion in the *ras2Δ* mutant, again hinting at a Ras/PKA-independent pathway that provides an additional benefit. The *msn2/4Δ* mutant on the other hand shows a significantly different diffusion phenotype only in the *HS SC +AntA* condition, where the diffusion is drastically lower, and the confinement is strongly increased compared to the wild type (**Figure 4F–H**). In *SC* and *SC +AntA*, the diffusion in *msn2/4Δ* is not significantly different than in the wild type. These results imply that in 1 hour of glucose starvation or respiration-deficient glucose starvation, the activation of *Msn2/4* is not strictly crucial for modulating the biophysical properties of the cytoplasm, but when Msn2/4 activation occurs, such as in the HS condition or in the *ras2Δ* strain, the cytoplasmic properties are modulated and the diffusion drop in the starvation conditions is partially counteracted.

This data therefore supports the hypothesis that activation of the ESR through Msn2/4 provides a beneficial adaptation of the cytoplasmic properties which correlates with a benefit to cell survival.

## Discussion

In this work, we demonstrate that the biophysical properties of the budding yeast cytoplasm are highly adaptive and that proper cytoplasmic reorganization correlates with viability in different quiescence conditions across genetic backgrounds. Preconditioning cells with a pre-HS before respiration-deficient starvation primes them for survival without restoring ATP levels (**Figure 1B and 1C**), indicating that energy availability alone does not dictate cell fate in quiescence. Instead, a stress-induced proteomic adaptation mediated in part by the Msn2/4-controlled ESR modulates cytoplasmic diffusion and plays a crucial role in long-term survival by maintaining cell viability and allowing re-entry into the cell division cycle (**Figures 3 and 4**).

Intriguingly, the HS-induced proteome differs from the adaptation that occurs during prolonged glucose starvation alone, regarding both global proteome changes as well as ESR-specific changes (**Figures 3G, S3A-S3C**). This is in line with previously studied stress-specific ESR regulation by Msn2/4 (Gasch et al., 2000). Moreover, some ESR genes that do not strictly require Msn2/4 for induction are regulated not only by Msn2/4 but also Hsf1 (Amorós and Estruch, 2001; Yamamoto et al., 2008). The expression of the small heat shock protein Hsp26, for example, requires Msn2/4 in glucose starvation, but in HS the Hsf1-mediated induction can compensate for a lack of Msn2/4 (Amorós and Estruch, 2001). The Hsf1-mediated stress response might explain the incomplete survival defect of the *msn2/4Δ* mutant in respiration-deficient starvation as well as the residual HS benefit that occurs in the *msn2/4Δ* mutant with regard to survival and cytoplasmic diffusion. Since *HSF1* itself is essential, its impact on survival cannot be detected in the SATAY screen. Additionally, many transcriptional Hsf1 targets are redundant and cannot score highly in the SATAY screen, even if they are contributing to the HS-mediated benefit. The functional redundancy of Msn2/4 and Hsf1 targets might also provide an explanation for the lack of overlap between the SATAY hits and the proteomic changes (**Figure S5A**). Furthermore, the signaling genes that scored highly in the SATAY screen are often regulated through their activity rather than their abundance. Plausibly, their downstream target proteins are differentially expressed, whereas disrupting the regulators themselves leads to strong survival phenotypes. Similar observations have been made in other stress conditions, where fitness-relevant genes showed little overlap with genes with upregulated expression (Birrell et al., 2002; Giaever et al., 2002).

Our in-depth analysis shows that yeast cells undergo significant cytoplasmic reorganization during glucose starvation, characterized by increased macromolecular crowding, reduced diffusion rates, and more frequent confined particle movement within smaller compartments (**Figures 1D–H and 2**). FIB-SEM imaging revealed a 28% higher macromolecular density in the cytoplasm of glucose-starved cells (**Figure 2A**), consistent with prior findings (Joyner et al., 2016) and similar observations by cryoelectron tomography in energy-depleted cells (Marini et al., 2020). In logarithmically growing cells, ribosomes that are organized into polysomes have been shown to contribute significantly to crowding and to influence the diffusion of probes in the size range of our 40-nm GEM particles (Gade et al., 2024; Xie et al., 2024),. Upon glucose starvation, polysomes collapse into monosomes (Ashe et al., 2000), yet ribosomes are expected to continue to contribute substantially to crowding and to modulate GEM diffusion. Recent studies have indicated that the biophysical state of the cytoplasm, including crowding, modulates the speed of biochemical reactions such as mRNA translation and protein degradation (Chen et al., 2024; Sareen et al., 2025). Additional cytoplasmic rearrangements occur upon energy depletion, including filament formation or the formation of biomolecular condensates (Narayanaswamy et al., 2009; Noree et al., 2010; Petrovska et al., 2014; Prouteau et al., 2017; Riback et al., 2017; Saad et al., 2017; Marini et al., 2020; Stoddard et al., 2020; Cereghetti et al., 2021) highlighting a broader reorganization of the cytoplasm that could further affect the diffusion of macromolecules. In turn, the amount of crowding might also influence the propensity to form condensates and filaments (Heidenreich et al., 2020).

The correlation between diffusion and survival outcomes across different genetic backgrounds (*ras2Δ* and *msn2/4Δ* mutants) underscores a potential functional relevance of cytoplasmic organization for cell viability and aging. Adapting cytoplasmic organization could be used to balance the need for slower metabolic processes in quiescence and the minimal diffusion that is essential for cellular processes and therefore survival. Of note, diploid yeast cells undergo meiosis upon glucose starvation, and then regulate cytoplasmic diffusion during spore dormancy, a protective quiescent state enabling very long-term survival even under hostile conditions (Plante et al., 2023; Sakai et al., 2024). By contrast, extremely low diffusion, as seen in the absence of respiration in glucose starvation, might irreversibly impair cellular functions due to limited movement of proteins and RNA in and out of certain compartments, cellular territories and condensates. Such a model is supported by our analysis of how lower diffusion is the result of confined spaces becoming more prevalent, creating compartments that can trap macromolecules. This state of low diffusion might also enhance protein aggregation or preclude the refolding of damaged proteins, which could disrupt proteostasis and lead to toxic aggregates that are a hallmark of the aging process (López-Otín et al., 2013). Together, these effects on proteins, RNA and other macromolecules likely lead to loss of cell fitness and irreversible arrest of the cells, preventing their reentry into the cell division cycle.

Proteomic analysis, survival assays and diffusion measurements suggest that PKA inhibition and Msn2/4 activation lead to stress response induction and the expression of genes required for proteostasis and protein folding, which is critical to ensure long-term survival in quiescence. We speculate that this, along with additional effects, prevents or reverts the extremely low diffusion state of respiration-deficient, starved cells. Protective proteins such as Hsps, which are regulated by both Msn2/4 and Hsf1, might contribute to the HS rescue effect, since they are clearly induced in *HS SC +AntA* and absent in *SC +AntA*. The large redundancy in the heat shock response might explain why they do not score in our SATAY screen. However, they are not strongly induced in *SC*, which suggests that cells do not strictly rely on them to maintain minimal diffusion. The mechanism behind the cytoplasmic reorganization and its beneficial effect on quiescence survival and chronological aging therefore remains to be elucidated further.

## Acknowledgments

The authors would like to thank the members of the Weis lab for discussions and comments on the manuscript. We thank Claudio De Virgilio for reagents and discussions. We would also like to acknowledge ScopeM for their support & assistance in this work. We are very grateful to Annett Neuner and Elmar Schiebel at Heidelberg University for their support in collecting yeast samples for FIB-SEM and providing the instruments for cell synchronization by elutriation.

This work was supported by grants from the European Research Council to P.P. (866004) from the Wellcome Trust to B.K. (214291/Z/18/Z) and from the Swiss National Science Foundation to K.W. (TMAG-3_209354 and CRSII5_193740).

## Author contribution statement

Conceptualization: LK, CAW, KW

Data curation: LK, CAW, PAG, JSF, CD, AM, PR, BK, KW

Formal analysis: LK, CAW, PAG, JSF, CD, AM

Investigation: LK, CAW, CD, AM, PR

Visualization: LK, CAW, JSF

Methodology: PAG

Software: PAG, JSF

Supervision: PAG, BK, KW

Writing – original draft: LK, PAG, JSF

Writing – review & editing: LK, CAW, PAG, JSF, CD, SK, BK, KW

Resources: PR, PP, BK, KW

Funding acquisition: PP, BK, KW

Project administration: KW

## Supplementary Material

**Supplemental Tables S1-S3**: List of generated and used yeast strains, plasmids and oligonucleotides.

**Supplemental File 1**: Heatmap of standard scores (z-scores) across all conditions for proteins with abundance changes greater than 2 z-scores in any condition relative to non-stressed cells.

## Materials and Methods

### Strain generation

All strains used in this study are created from the W303 background. *Saccharomyces cerevisiae* strains were constructed using standard yeast genetic techniques either by transformation of a linearized plasmid or of a PCR amplification product with homology to the target site according to Longtine et al. (1998). Transformation of *Saccharomyces cerevisiae* cells was performed according to Gietz and Woods (2006). Used strains, plasmids and primers are listed in Supplemental Tables S1–S3.

### Yeast cultivation

Cells were grown at 30 °C in synthetic complete (SCD) medium with glucose (2% w/v glucose (Sigma), 6.7g/l yeast nitrogen base (Difco), essential amino acids (Sigma, Merk), pH 5.0) overnight. Cultures were diluted in fresh SCD to an OD600 of 0.2 and grown into logarithmic phase (OD600 0.4-0.9).

### Glucose starvation

Logarithmically growing cells were washed three times into medium lacking glucose (SC), either by pelleting the cells and replacing the medium or by directly replacing the medium of cells in microscopy wells.

### Spotting assays

Cells were spotted according to the OD600 measured after washing into starvation medium. Starting at an OD600 of 0.2, cells were serially diluted 1:5 three times and spotted onto YPD plates, which were incubated for 2 days at 30 °C. YPD medium consisted of 20 g/l bacto agar (BD Biosciences), 20 g/l bacto peptone (BD Biosciences), 10 g/l yeast extract (BD Biosciences), 2% w/v glucose (Sigma), and 80 mg/l Adenine Hemisulphate (Sigma). The YPD plates were scanned with an Epson Scanner. The scan images were processed in Fiji (Schindelin et al., 2012) (ImageJ 1.54f).

### Heat shock treatment

Exponentially growing cells were incubated in a 42 °C water bath for 30 min. Correct culture temperature was checked with an additional tube containing water. Cells were then recovered in a rotating wheel at 30 °C for 30 min. Subsequently, cells were treated according to the specification within the experiment.

### Drug treatments

Antimycin A (AntA, Sigma), dissolved in DMSO, was used at a final concentration of 10 μM in all experiments. Antimycin A was added to the glucose-free medium (SC) prior to washing the cells with it. Cycloheximide (CHX, Sigma) was dissolved in water and used at a final concentration of 10 μg/mL in all experiments. Cycloheximide was added to logarithmically growing cells for 5 minutes, then the cells were heat-shocked (42 °C, 30 min) and recovered (30 °C, 30 min) with the Cycloheximide present. Upon washing the cells into glucose-free medium, the Cycloheximide was washed out and not re-added. 1-naphthylmethyl PP1 (1NM-PP1, Sigma), dissolved in DMSO, was used at a final concentration of 5 μM in all experiments. 1NM-PP1 was added to logarithmically growing cells for 3 hours, then the cells were either kept at 30 °C for 1 hour or heat-shocked (42 °C, 30 min) and recovered (30 °C, 30 min) with the 1NM-PP1 present. Upon washing the cells into glucose-free medium, the 1NM-PP1 was washed out and not re-added.

### ATP measurements

ATP measurements were done according to Seo et al. (2017) with minor modifications. Cells were pelleted and resuspended in 750 μl 90% acetone. Normalization was done according to OD600. The cells were then incubated at 90°C for approx. 10 min to evaporate the acetone, when about 50 μl of solution remained. 450 μl of buffer (10mM Tris pH 8.0, 1mM EDTA) was added to the solution and ATP was measured with an ATP Determination Kit (Thermo Scientific) on a CLARIOstar microplate reader (BMG).

### HILO Imaging of 40 nm GEMs

384-well plates (Matrical) were coated with concanavalin A (Sigma, C201) for a few minutes. 30 μl of yeast cell suspension after respective treatments were plated in each well. The plate was centrifuged at 100 g for 1 min and imaged at 30 °C. Strains expressing 40 nm GEMs were imaged at 50% laser power using 488 nm (80 mW) laser excitation on a Nikon N-STORM microscope in HILO (Highly Inclined and Laminated Optical) mode. Imaging was performed with an SR Apochromat TIRF 100X (NA=1.49) oil immersion objective and sCMOS camera: Hamamatsu Orca Flash 4 v3 (6.5 x 6.5 μm² pixel size). We used an additional demagnification lens system with a 2.4X de-magnifying power. The effective pixel size is 160 nm (6.5 μm/100X*2.4X). Images were acquired at ∼33 frames/sec for 21-30 seconds using NIS Elements Advanced research software. Images were processed using Fiji (Schindelin et al., 2012) (ImageJ 1.54f).

### Single particle localization and tracking

Tracking of GEM particles was performed with the TrackMate 7 (Ershov et al., 2022) plugin of Fiji (Schindelin et al., 2012) (ImageJ 1.54f). For the localization step, the LoG detector with sub-pixel localization enabled was used and for the tracking step, the Simple LAP tracker. Spot radius = 0.225 microns, Quality threshold = 100.0. Maximum number of frames to close the gap = 3. Maximum distance for closing a gap = 0.7 microns. Maximum jump from one frame to the next = 0.32 microns. A minimum track length of 4 frames was used.

### Trajectory analysis: Trajectory classification

All generated trajectories are analyzed with a custom written Matlab(R2021) pipeline which uses the @msdanalyzer library (Tarantino et al., 2014). The codes are available here: https://github.com/PabloAu/Single-Molecule-Tracking-Analysis/tree/master/V2. First, the Mean Squared Displacement (MSD) is computed for an individual trajectory and a motion type classification is performed, to split the trajectories into confined and unconfined subgroups (Gómez-García et al., 2021). This classification is done by fitting a power law function to an individual Time-averaged MSD (T-MSD) curve and obtaining the anomalous exponent coefficient α:

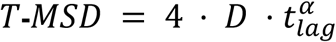

where D is the diffusion coefficient, tlag is the time lag between the different time points of the track, and α is the anomalous exponent coefficient. Trajectories with α≤0.8 were considered as confined and trajectories with α>0.8 as unconfined. The percentage belonging to each population is shown in several figures.

### Trajectory analysis: Diffusion Coefficients

Diffusion coefficients were obtained in a population-based fashion, by fitting the first 4 points of the Time-Ensemble MSD (TE-MSD) curves with a linear function:

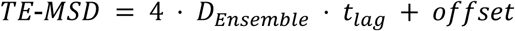

This is more accurate than looking at the individual Time MSD curves since the number of jumps to compute the TE-MSD curves is much higher (Gómez-García et al., 2021). The ensemble diffusion coefficients (*D* = *D_Ensemble_*) are shown across the different figures.

### Trajectory analysis: Radius of Confinement

The radius of confinement was obtained only for the subgroup of confined trajectories. It quantifies the degree of confinement by estimating the radius of the potential volume explored by the particle. It was measured by fitting a circle-confined diffusion model to the TE-MSD (ensemble of all trajectories) (Wieser and Schütz, 2008).

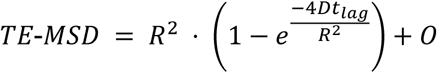

where *R* is the radius of confinement and *D* is the diffusion coefficient at short timescales. *O* is an offset value that comes from the localization precision limit inherent to localization-based microscopy methods.

### Trajectory analysis: Track Total Intensity

For each individual particle trajectory, the frame with the highest signal intensity of the localized particle was determined and the sum of the pixel intensities of the particle in that frame was calculated as the “Track Total Intensity”.

In fixed cells, the GEM intensities were comparable in all conditions (**Figure S2C**). All GEM intensity histograms show a single, bell-shaped distribution of intensities with no indication of several GEM particles aggregating into brighter foci.

### Cell synchronization by elutriation for FIB-SEM

Elutriation of small G1 cells (unbudded) was performed using a centrifugal elutriation system (Beckman Coulter) in a standard 4 ml chamber. The elutriator was used according to the manufacturer’s protocol. The centrifuge was constantly running at 2500 rpm, and cells were flushed into the chamber at 7 ml/min. Once the chamber was filled, cells were eluted at 8 ml/min. The eluted cells were immediately washed into SCD or SC medium. Synchrony of the population was calculated to be 91%. The control cells in SCD were kept at 30 °C for 10-15 min after the washes, and then high pressure frozen, while the SC cells were incubated at 30 °C for 30 min after the washes and then frozen for further processing with FIB-SEM.

### FIB-SEM

All samples were high pressure frozen using a HPM010 system in Abra Fluid. Freeze substitution was performed with 1% OsO4, 0.2% uranyl acetate, and 5% water in dry acetone according to the following protocol: the sample was kept at -90 °C for 64 h, the temperature was then increased in 5 °C/h steps to -30 °C, where incubation was continued for 4 h. Temperature was then raised in 5°C/h steps to 20 °C, where the sample remained at for 5 h. Following the freeze substitution, the samples were rinsed three times in acetone and incubated for 20 min at room temperature in 0.1% thiocarbohydrazide and 10% water. After another threefold rinse with acetone, samples were incubated in 2% OsO4 in acetone using a microwave (BioWave, TedPella). Samples were rinsed again 4 times in acetone, before proceeding with the infiltration in Hard Plus resin (EMS) using the microwave (protocol: 2x10% – 2x25% – 2x50% + 60 min RT – 2x75% + 60 min RT – 2x90% + 60 min RT – 2x100% + 16 h RT – 2x100% – polymerization at 60°C for 72 h). The samples were then trimmed and mounted on an aluminum stub for SEM. The samples were imaged in a Zeiss CrossBeam 550 FIB-SEM (Carl Zeiss Microscopy) using the Atlas Nanotomography workflow (Fibics). FIB milling was achieved at 30 kV with a current of 1.5 nA. SEM imaging was performed with 1.5 kV acceleration voltage and 700 pA current of the electron beam and an ESB detector (grid 1100V). The voxel size of the acquisition was 5 nm isotropic. Processing of images was performed in Fiji (Schindelin et al., 2012) (ImageJ 1.54f), using a linear SIFT alignment plugin based on Lowe (2004). Ribosome counting was done on at least 3 volumes per condition, and the regions chosen did not contain any organelles or structures larger than single molecules. Moving through the 3D slices, the electron dense spots were counted manually.

### Hyperosmotic stress treatment

Cells were grown exponentially in SCD medium. SCD medium containing 2X of the desired NaCl (Sigma) concentration was added 1:1 directly to the cell culture in the microscopy well. Cells were imaged after 5 min of SCD +NaCl addition.

### Cell volume determination

Images obtained for GEM particle tracking were processed in Fiji (Schindelin et al., 2012) (ImageJ 1.54f) by creating an average projection of the 700-frame movie. To this average projection, CellPose (TrackMate 7 (Ershov et al., 2022)) was applied (initial radius: 4.0 μm, min. circularity = 0.81, min. radius = 0.6 μm, max. radius = 6.0 μm) in order to obtain the ROI and the radius [μm] for each cell. Assuming a spherical cell shape, the radius (r) was used to calculate the cell volume = (4/3 * r * π)^3^. In each replicate, the average cell volume was determined based on the radii determined from at least 400 cells.

### Proteomics: Cell Harvest and Lysis

Yeast cell pellets were resuspended in lysis buffer consisting of 20 mM HEPES (Sigma, H4034), 150 mM KCl (Merck, 1049360250), and 10 mM MgCl₂ (Sigma, M2670), adjusted to pH 7.5, supplemented with 1× Roche Complete Protease Inhibitor EDTA-free (Sigma-Aldrich, 11873580001). The suspension was transferred to screw-cap microcentrifuge tubes, and an equal volume of glass beads was added. Cells were lysed at 4 °C using a benchtop homogenizer (Fastprep-24 5G, MP Biomedicals) for eight cycles of 30 s beating at 5.5 Hz, with 200 s cooling intervals between cycles. Tubes were perforated on both top and bottom using a heated needle, placed onto 1.5 ml collection tubes, and centrifuged at 1000 g for 120 s to recover the lysate.

### Proteomics: Protein Concentration Measurement

Protein concentration in the lysates was determined using the bicinchoninic acid assay (BCA; Pierce BCA Protein Assay Kit, Thermo Fisher Scientific, 23225). Samples were adjusted to 2 µg/µl, corresponding to a total of 100 µg protein in 50 µl.

### Proteomics: Tryptic Digest

Disulfide bonds were reduced by adding tris-(2-carboxyethyl)phosphine (TCEP; Sigma, C4706) from a 200 mM stock to a final concentration of 10 mM, followed by incubation at 37 °C for 40 min with shaking at 800 rpm. Alkylation was performed by adding freshly prepared 1 M iodoacetamide (Sigma, I1149) to a final concentration of 40 mM, with incubation at room temperature for 25 min in the dark without shaking. Samples were diluted 1:5 with 100 mM ammonium bicarbonate (Sigma-Aldrich, A6141) to yield a final sodium deoxycholate (DOC) concentration of 1% (w/v). Proteolysis was initiated by adding 1 µl lysyl endopeptidase R (FUJIFILM Wako Pure Chemical Corporation, 129-02541) and 2 µl trypsin (0.5 mg/ml; Promega, V511C). Digestion proceeded overnight at 37 °C with shaking at 350 rpm. The reaction was quenched by adding 100% formic acid (Carl Roth GmbH) to a final concentration of 3% (v/v), reducing the pH below 2. DOC was removed by centrifugation at 1200 g for 2 min.

### Proteomics: C18 Cleanup and Sample Preparation

Peptide cleanup was performed using a 96-well C18 spin column plate (Nest Group, S8VL) connected to a vacuum manifold. Columns were sequentially conditioned with 100 µl methanol (Fisher Scientific, 15631400), 50 µl buffer B (50% acetonitrile [ACN], Fisher Scientific, A955-212; 0.1% formic acid, Carl Roth GmbH), and 3 × 100 µl buffer A (5% ACN, 0.1% formic acid). Peptide samples were loaded, washed with 3 × 100 µl buffer A, and eluted with 3 × 100 µl buffer B. Eluates were dried using a vacuum centrifuge and resuspended in buffer A to 2 mg/ml peptide concentration. Indexed Retention Time (iRT) peptides (10×; Biognosys, Pp-2005) were spiked into each sample at a 1:30 dilution.

### Proteomics: Data Acquisition and Analysis

The acquired DIA data was searched in Spectronaut v18.6 (Biognosys) using the directDIA algorithm. Non-tryptic peptides or peptides with more than two missed cleavages were excluded from the analysis. Carbamidomethylation was used as a fixed modification and methionine oxidation was set as variable modification. Spectra were searched against the SGD protein database (downloaded on 13.10.2015, 6713 entries) using a 1% FDR control at peptide and protein level. For the chromatogram extraction, default settings were used except “cross-run normalization” was excluded. The ion intensities at the fragment level were then further analyzed in R. In brief, low-quality fragment ions were excluded based on the Spectronaut ‘F.ExcludedFromQuantification’ flag and only proteotypic precursor ions were retained. For each precursor the remaining fragment ions were summed as the respective precursor intensity. To filter out low quality precursor ions, the median protein intensity was calculated from all precursors and for each protein, the precursor ions with the highest 25% mean square error from the median intensity across all samples were filtered out. Protein intensities were then calculated using the maxLFQ algorithm (Cox et al., 2014). Proteins that were not covered by at least 5 precursors in all four replicates of minimum one condition were not considered for further analysis (3308 proteins remaining). Intensities for protein missing in a replicate were imputed as follows. A normal distribution was fitted to the bottom 1% protein intensities and values were randomly sampled from this distribution (in total 118506 non-imputed, 582 imputed protein intensities). All protein intensities were then median normalized across samples and the mean of all four replicates was taken per condition. Significance in volcano plots was calculated using a two-sided unpaired Student’s t-test and resulting p-values were adjusted for multiple testing using the Benjamini-Hochberg correction. For gene ontology (GO) analysis, the significance of the GO-terms of the highest or lowest 10% differentially abundant proteins between two conditions was assessed using a hyper-geometric test and resulting p-values were adjusted for multiple comparisons using the Benjamini-Hochberg correction. Significant but highly redundant GO-terms were manually excluded and not shown in Figure 3F. For Figure S3A, all GO terms that were significantly enriched with a q-value higher than 3 in any stress condition as compared to non-stressed cells were plotted.

### SATAY Screen: Experimental Design

For the screen, the cultured SATAY library was split into three parts. The first was treated with a pre-heat shock (30 min at 42 °C), recovery (30 min at 30 °C) and subsequent respiration-inhibited (AntA) glucose starvation for 4 days (HA4d). The two additional samples were used as controls. One was a control for short-term effects of respiration-inhibited glucose starvation (A1), where cells were starved for one hour in the presence of Antimycin A. The second control was pre-heat shocked, left to recover and then starved for glucose for one hour in the presence of Antimycin A (HA1) to control for effects of the heat shock prior to starvation. After treatment, all cultures were harvested and re-grown in complete medium (YPD) to assess which mutants survived and were able to proliferate after the stress.

### SATAY Screen: Sample Preparation and Harvest

W303 wild-type cells were transformed with pBK549 and grown to an OD600 of 4-6 in SC medium lacking uracil with 0.2% glucose (Sigma, G8270) and 2% raffinose (Fluka, 83400) added. To induce transposition, cells were grown in SC medium lacking uracil and with 2% galactose (Sigma, G5388) added for 50-52 h. The number of transposed cells was evaluated by plating cells onto SCD plates lacking adenine. The SATAY library was then concentrated, 10% DMSO was added, and the mixture was put at -20 °C overnight and then stored at -80 °C. Viability and amount of transposed (ADE+) cells was again assessed in the frozen library, before inoculating 50 million viable transposed cells into synthetic complete medium lacking adenine (SCD –ADE). This pre-culture was grown for four population doublings, and diluted whenever necessary to maintain the culture in exponential growth. The culture was then split into three samples. Two of these samples (HA1, HA4d) were incubated for 30 min at 42 °C in a water bath with a shaker. After incubation, the samples were cooled down 5 min in a shaker at 25 °C, as well as for 25 min at 30 °C shaking. The third sample (A1) was maintained during this 1 h at 30 °C in exponential growth phase. All three samples were then washed three times into SC medium with 10 µM Antimycin A added (see glucose starvation protocol for more details). All cultures were then incubated at 30 °C on a shaker. The HA1 and A1 samples were harvested after 1 h of starvation, while the HA4d sample was harvested after four days in respiration-deficient starvation. For the harvest, cells were inoculated at an OD600 of 0.25 in 500 ml YPD and grown to saturation at 30 °C shaking. Cells were centrifuged and 500 mg of the cell pellet were used for DNA extraction. DNA extraction and library sequencing were performed according to Michel et al. (2017).

### SATAY Screen: Analysis and Plotting

For each gene in the three sequenced libraries (A1, HA1, HA4d), the number of transposons per gene (#tn_per_gene) and the number of transposon reads per gene (#read_per_gene) were determined, and an arbitrary, small number was added to each gene (+5 to #tn_per_gene, +25 to #read_per_gene) to reduce noise. For normalization, the #tn_per_gene of each gene was divided by the total number of transposons or reads sequenced in the given library, and the #read_per_gene of each gene was divided by the total number reads sequenced in the given library. The libraries were then assigned to a test (HA4d) or control (A1, HA1) set. For each gene, and individually for the #tn_per_gene or #read_per_gene, the test set was divided by the average of the control sets, to obtain the fold change, which is plotted on the Volcano plot’s x-axis. We then determine the -log10 of the p-value of a Student’s t-test between the test and control set for each gene, which is plotted in the Volcano plot’s y-axis. This results in two Volcano plots, one based on the number of transposons per gene (#tn_per_gene) and one based on the number of transposon reads per gene (#read_per_gene). To obtain one meaningful Volcano plot, we merged the two plots, taking the data point with the lower p-value for each gene. In general, the genes that are enriched in transposons in the test set (positive fold change), have a lower p-value in the #read_per_gene data set, since they have a fitness increase and divide more, which leads to more transposon reads being sequenced. On the other hand, genes with few transposon insertions have a lower p-value in the #tn_per_gene data set. In this mixed Volcano plot, the negative fold change side mostly contains #tn_per_gene data and the positive fold change side mostly the #read_per_gene data.

## Supplemental Figures

**Supplemental Figure S1:**
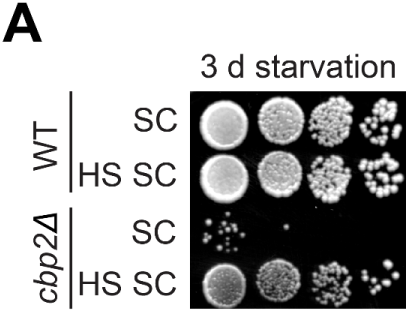
Major survival defect in respiration-deficient glucose starvation is rescued by preparatory heat shock. **A)** Survival assay of wild-type (WT) and *cbp2Δ* cells treated with the indicated conditions and starved for 3 days prior to spotting and re-growth on YPD plates.

**Supplemental Figure S2:**
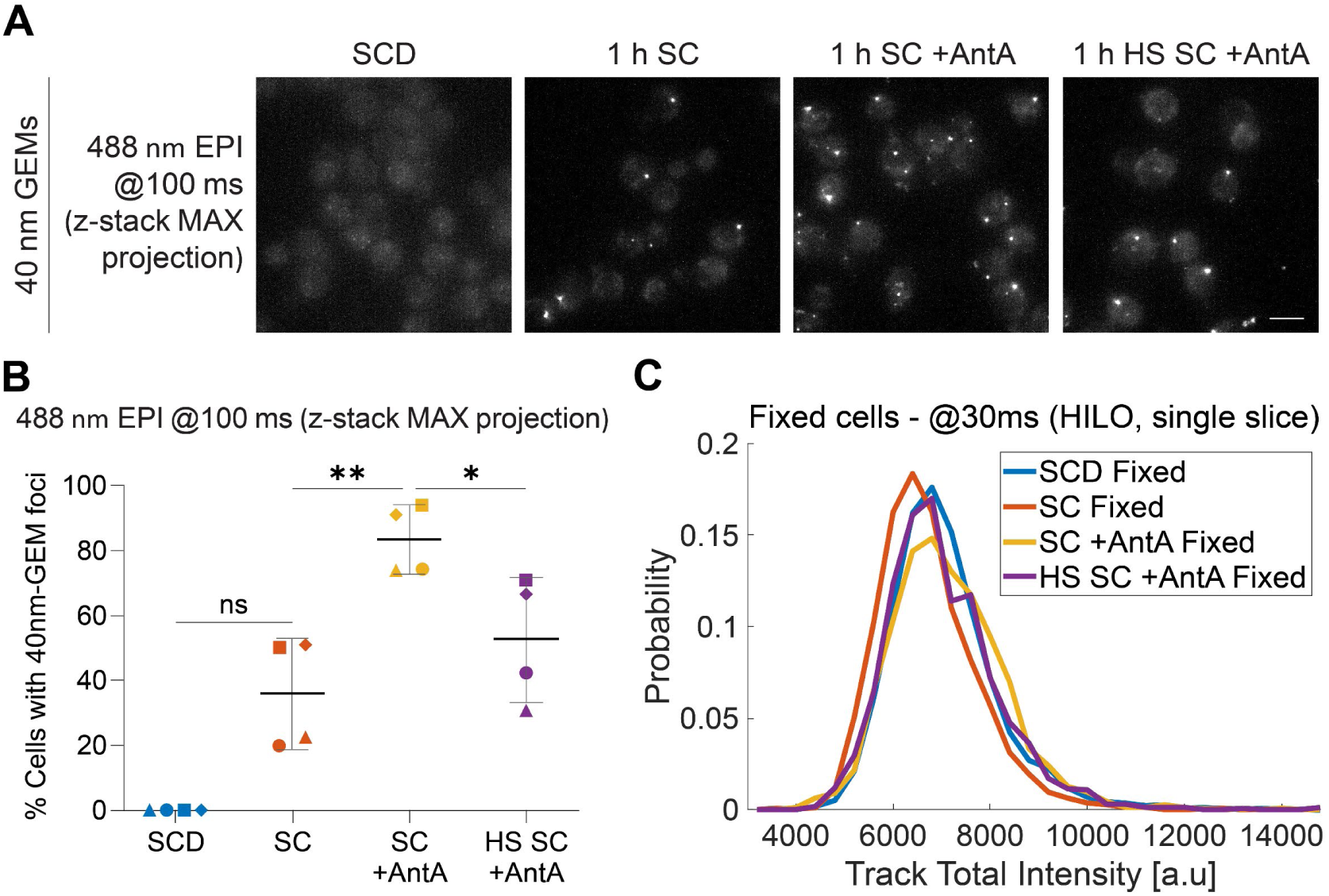
Quiescence causes confinement of GEM particles, not aggregation. **A)** GEM-expressing cells in indicated conditions were imaged at 100 ms exposure time (10 FPS, 488 nm EPI, z stack projection). **B)** Quantification of percentage of cells imaged at 100 ms exposure time (10 FPS, 488 nm EPI, z stack projection) with visible GEM foci (n = 4 independent replicates, mean±SD). One-way ANOVA with Tukey’s multiple comparisons test (^ns^p > 0.05, *p ≤ 0.05, **p ≤ 0.01). **D)** Line histogram showing the distribution of the Track Total Intensities determined from GEM trajectories in fixed cells in SCD, SC, SC +AntA and HS SC +AntA (n = 3 independent replicates, one replicate is shown). The cells were imaged at 30 ms exposure time (33.3 FPS, 488 nm HILO, single slice).

**Supplemental Figure S3:**
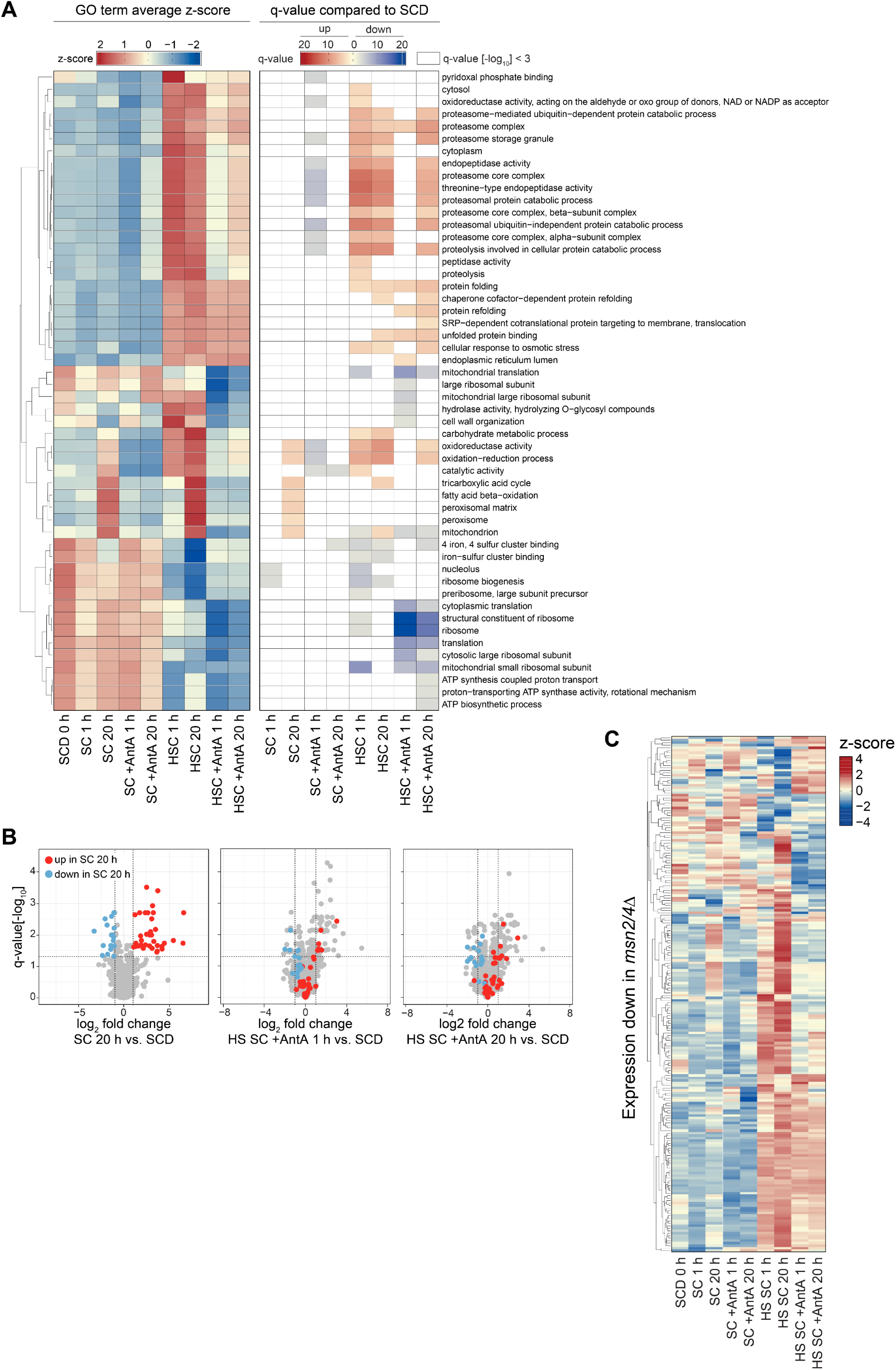
Proteome analysis reveals distinct adaptation in glucose starvation and heat shock conditions. **A)** GO terms that are significantly up or down in any condition as compared to non-stressed cells. **Left**: Heatmap of standard scores (z-score) of the mean abundance of all proteins associated with the indicated GO term. **Right:** Heatmap of q-values calculated by testing the 10% highest and lowest abundant proteins for significance using a hyper-geometric test and adjusting the resulting p-values for multiple comparisons using the Benjamini-Hochberg correction. **B)** Proteins that showed up- or downregulated abundance in SC 20 h compared to SCD are highlighted in the comparison plots of protein changes between HS SC +AntA vs. SCD. **C)** Heatmap of standard scores (z-score) across all conditions in this study for proteins that were downregulated in the *msn2/4Δ* mutant (Kuang et al., 2017). Every row corresponds to one protein.

**Supplemental Figure S4:**
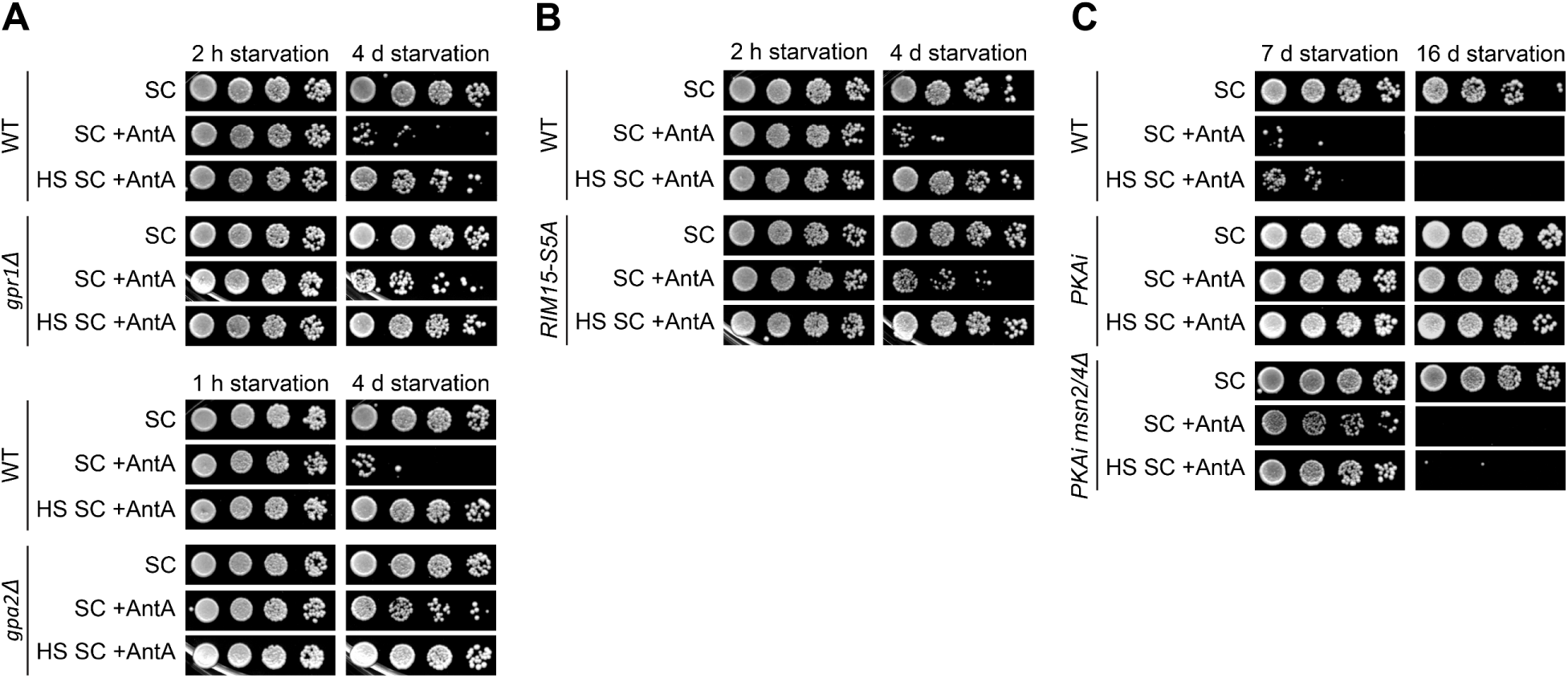
Glucose signaling and stress response activation impact quiescence survival. **A-C**) Survival assays of wild-type (WT) and mutant cells treated as indicated and starved for the indicated time were spotted onto YPD plates. Regrowth ability is used to qualitatively assess survival. The constitutively active RIM15 mutant (RIM15-S5A (Reinders et al., 1998)) was overexpressed using an estradiol-inducible promoter.

**Supplemental Figure S5:**
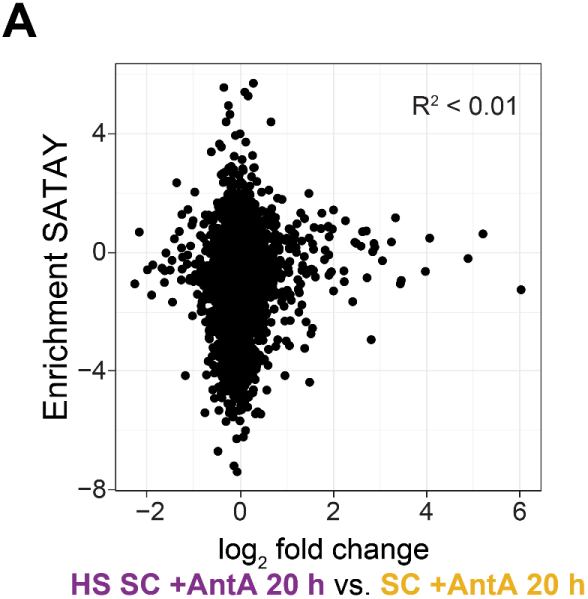
The SATAY screen and proteomics show poor correlation. **A**) Correlation between the enrichment in the SATAY screen and the log2 fold difference in protein abundance comparing respiration-deficient 20 h glucose starved cells with (HS SC +AntA) or without (SC +AntA) a preparatory-heat shock. Each point corresponds to one protein. Coefficient of determination R^2^ < 0.01. z-score: Each protein intensity was standardized by subtracting the mean protein intensity across all conditions and dividing by the standard deviation.

