## Supplementary figures and images for "The environmental stress response controls the biophysical properties of the cytoplasm and is critical for survival in quiescence"

### Supplemental File 1

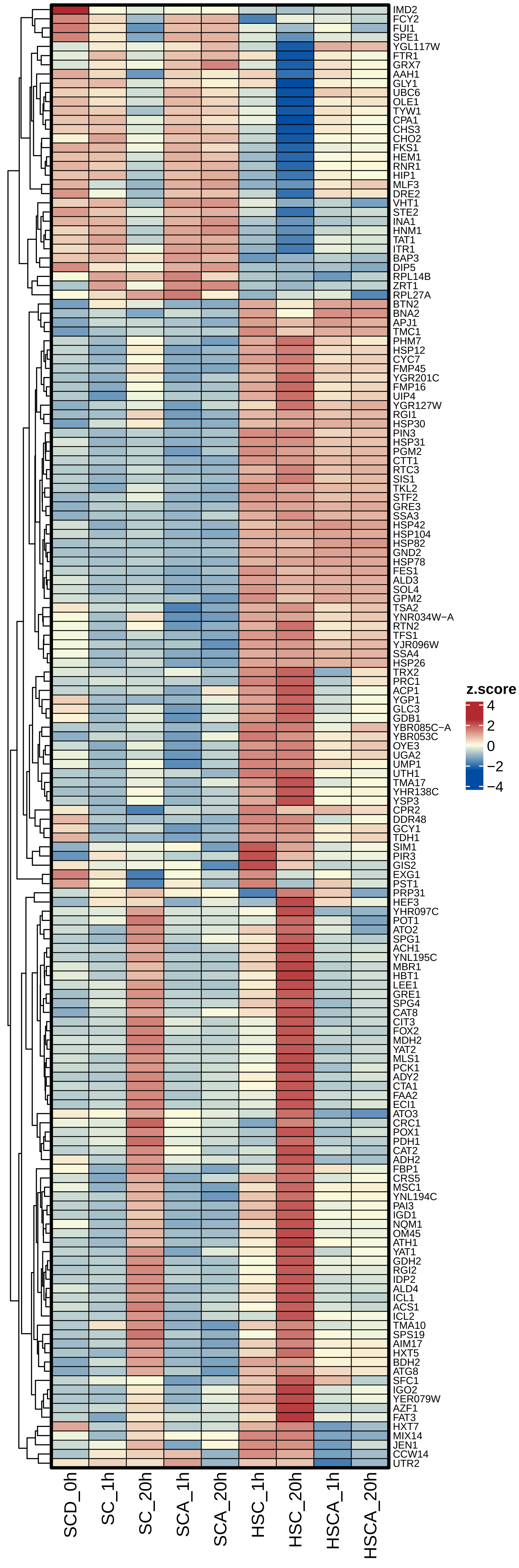
